# A Ssd1 homolog impacts trehalose and chitin biosynthesis and contributes to virulence in *Aspergillus fumigatus*

**DOI:** 10.1101/594341

**Authors:** Arsa Thammahong, Sourabh Dhingra, Katherine M. Bultman, Joshua Kerkaert, Robert A. Cramer

## Abstract

Regulation of fungal cell wall biosynthesis is critical to maintain cell wall integrity in the face of dynamic fungal infection microenvironments. In this study, we observe that a yeast *ssd1* homolog, *ssdA,* in the filamentous fungus *Aspergillus fumigatus* is involved in trehalose and cell wall homeostasis. An *ssdA* null mutant strain exhibited an increase in trehalose levels and a reduction in colony growth rate. Over-expression of *ssdA* in contrast perturbed trehalose biosynthesis and reduced conidia germination rates. The *ssdA* null mutant strain was more resistant to cell wall perturbing agents while over-expression of *ssdA* promoted increased sensitivity. Over-expression of *ssdA* significantly increased chitin levels and both loss and over-expression of *ssdA* altered sub-cellular localization of the class V chitin synthase CsmA. Strikingly, over-expression of *ssdA* abolished adherence to abiotic surfaces and severely attenuated the virulence of *A. fumigatus* in a murine model of invasive pulmonary aspergillosis. In contrast, despite the severe *in vitro* fitness defects observed upon loss of *ssdA,* neither surface adherence or murine survival was impacted. In conclusion, *A. fumigatus* SsdA plays a critical role in cell wall homeostasis that alters fungal-host interactions.

**Importance:** Life threatening infections caused by the filamentous fungus *Aspergillus fumigatus* are increasing along with a rise in fungal strains resistant to contemporary antifungal therapies. The fungal cell wall and the associated carbohydrates required for its synthesis and maintenance are attractive drug targets given that many genes encoding proteins involved in cell wall biosynthesis and integrity are absent in humans. Importantly, genes and associated cell wall biosynthesis and homeostasis regulatory pathways remain to be fully defined in *A. fumigatus.* In this study, we identify SsdA, a model yeast Ssd1p homolog, as an important component of trehalose and fungal cell wall biosynthesis in *A. fumigatus* that consequently impacts fungal virulence in animal models of infection.

## Introduction

*Aspergillus fumigatus* is the most common filamentous fungus that causes a wide variety of human diseases ranging from allergic type diseases to acute invasive infections. Invasive aspergillosis (IA) in immune compromised hosts, e.g. patients with hematological malignancies and organ or stem cell transplant recipients is associated with high mortality (1). Antifungal drugs used to treat IA, e.g. voriconazole, amphotericin B, are associated with undesired side effects, detrimental drug-drug interactions, long therapeutic regimens, and persistence of poor patient outcomes (2–5). To compound the difficulty of treating these infections, the emergence of azole-resistant *Aspergillus* infections is increasing globally (6–28). In order to make therapeutic advances against these increasingly common and potentially drug resistant infections, new antifungal drugs are needed.

One existing antifungal drug target that has not been fully exploited is the fungal cell wall. Consisting mainly of polysaccharides including β-1,3-glucans, alpha-glucans, mannan, chitin, and galactosaminogalactan among others (29–38), the cell wall is a great antifungal drug target as evidenced by the development of the echinocandins that target β-1,3-glucan biosynthesis. Interestingly, the carbohydrates needed to generate β-1,3-glucan and chitin, i.e. glucose 6-phosphate and UDP-glucose, are also important substrates used to generate trehalose, a disaccharide sugar, that is important for fungal conidia germination, stress protection, cell wall homeostasis, and virulence (39,40). The canonical trehalose biosynthesis pathway in *A. fumigatus* consists of two enzymes, TpsA/B (trehalose-6-phosphate synthase) and OrlA (trehalose-6-phosphate phosphatase), and two regulatory-like subunits, TslA and TslB (39–41). Trehalose biosynthesis is also found in other organisms in addition to fungi including bacteria, plants, and insects, but is not found in humans (42).

We and others previously observed that proteins involved in trehalose biosynthesis impact cell wall homeostasis in *A. fumigatus* (39–41). Disruption of the trehalose-6-phosphate phosphatase, OrlA, leads to perturbations in cell wall integrity as observed by the increased sensitivity to the cell wall perturbing agents congo red, calcofluor white, and nikkomycin Z (40). Loss of OrlA attenuates virulence of *A. fumigatus* in chronic granulomatous disease (xCGD) and chemotherapeutic invasive pulmonary aspergillosis (IPA) murine models. In addition, a regulatory subunit of the trehalose biosynthesis pathway, TslA, is critical for trehalose production and cell wall homeostasis in part through regulation of a class V chitin synthase enzyme, ChsE/CsmA (41). Loss of TslA increased chitin production and altered the sub-cellular localization of CsmA (41). Pulldown assays with TslA as bait identified a physical interaction between TslA and CsmA as well as a putative *Saccharomyces cerevisiae* Ssd1 homolog, herein called SsdA.

Ssd1p is a pleiotropic RNA-binding protein (43, 44) that is important for chromosome stability at high temperature, vesicular trafficking, stress responses, and cell wall integrity in *S. cerevisiae* (45–49). Ssd1p genetically interacts with Pkc1p and Sit4p, which are important components of the cell wall integrity signaling pathway (48). It has been shown that *S. cerevisiae ssd1* null mutants isolated from both patients and plants are more virulent than wild type strains in a DBA/2 murine model (50). Similar to the *A. fumigatus tslA* null mutant, the cell wall composition of yeast *ssd1* null mutants contain more chitin and mannan with decreases in β-1,3-glucan and β-1,6-glucan (50). These cell wall changes lead to increased proinflammatory responses against *ssd1* mutant strains (50). In contrast to *S. cerevisiae,* the *ssdA/ssdA* null mutant in *C. albicans* has decreased virulence in the invasive systemic candidiasis murine model (51). In filamentous fungal pathogens, the function of Ssd1 homologs is less clear, but in the plant pathogen *Magnaporthe grisea, SSD1* is important for fungal colonization of rice leaves (52). The authors suggested that *SSD1* is essential for proper cell wall assembly leading to evasion of the host immune response (52). However, the mechanism(s) behind this phenotype is not well understood (52). In this study, we characterized a predicted Ssd1p homolog identified in *A. fumigatus* through its protein-protein interaction with the trehalose biosynthesis protein, TslA. Using a genetics approach, we observe that *A. fumigatus* SsdA is critical for cell wall biosynthesis, trehalose production, polarized growth, biofilm formation, and virulence of *A. fumigatus*. Our results support the known role of Ssd1 homologs in fungal cell wall biosynthesis and highlight the potential altered functions of *A. fumigatus ssdA* including those involved in trehalose biosynthesis, biofilm formation, and fungal virulence.

## Results

### SsdA regulates trehalose production and is required for *Aspergillus fumigatus* conidia germination and mycelium expansion

AFUB_010850 was identified in a mass-spectrometry based screen of proteins that interact with the trehalose biosynthesis regulatory protein, TslA (41). Protein domain analysis of AFUB_010850 revealed strong amino acid sequence similarity with the nucleic acid binding Interpro domain IPR012340 (5.0E^-109^) and the ribonuclease (RNB) PFAM domain PF00773 (4.4E^-88^). BLASTP analysis of the AFUB_010850 amino acid sequence against the *Saccharomyces cerevisiae* genome database revealed strong sequence similarity to the protein SSD1p. Consequently, reciprocal BLASTP analyses with SSD1p against the *A. fumigatus* genome suggested AFUB_010850 is likely an Ssd1p ortholog and hence we named AFUB_010850 *ssdA*. Given the previously identified roles of TslA in trehalose and cell wall homeostasis in *A. fumigatus,* and the known roles of *ssd1* homologs in fungal cell wall biosynthesis, we hypothesized that SsdA is an important mediator of trehalose production and cell wall homeostasis in *A. fumigatus*.

To test this hypothesis, we generated *ssdA* null mutant (Δ*ssdA)* and over-expression strains (OE:*ssdA)* (confirmed by PCR and Southern blot analyses). The *ssdA* null mutant was complemented with an *ssdA:GFP* allele. We next measured the trehalose content of both conidia and mycelia in the respective strains (53). We observed increased trehalose content in Δ*ssdA* conidia and mycelia and conversely reduced trehalose levels in OE:*ssdA* (*P=*0.0004, compared Δ*ssdA* to the wild type; *P*<0.0001 compared OE:*ssdA* to the wild type) (**Fig. 1A**). Reconstitution of Δ*ssdA* with *ssdA:GFP* restored wild-type trehalose levels. These data suggest that SsdA plays a role in regulating trehalose biosynthesis and/or levels in *A. fumigatus*.

**Fig 1.**
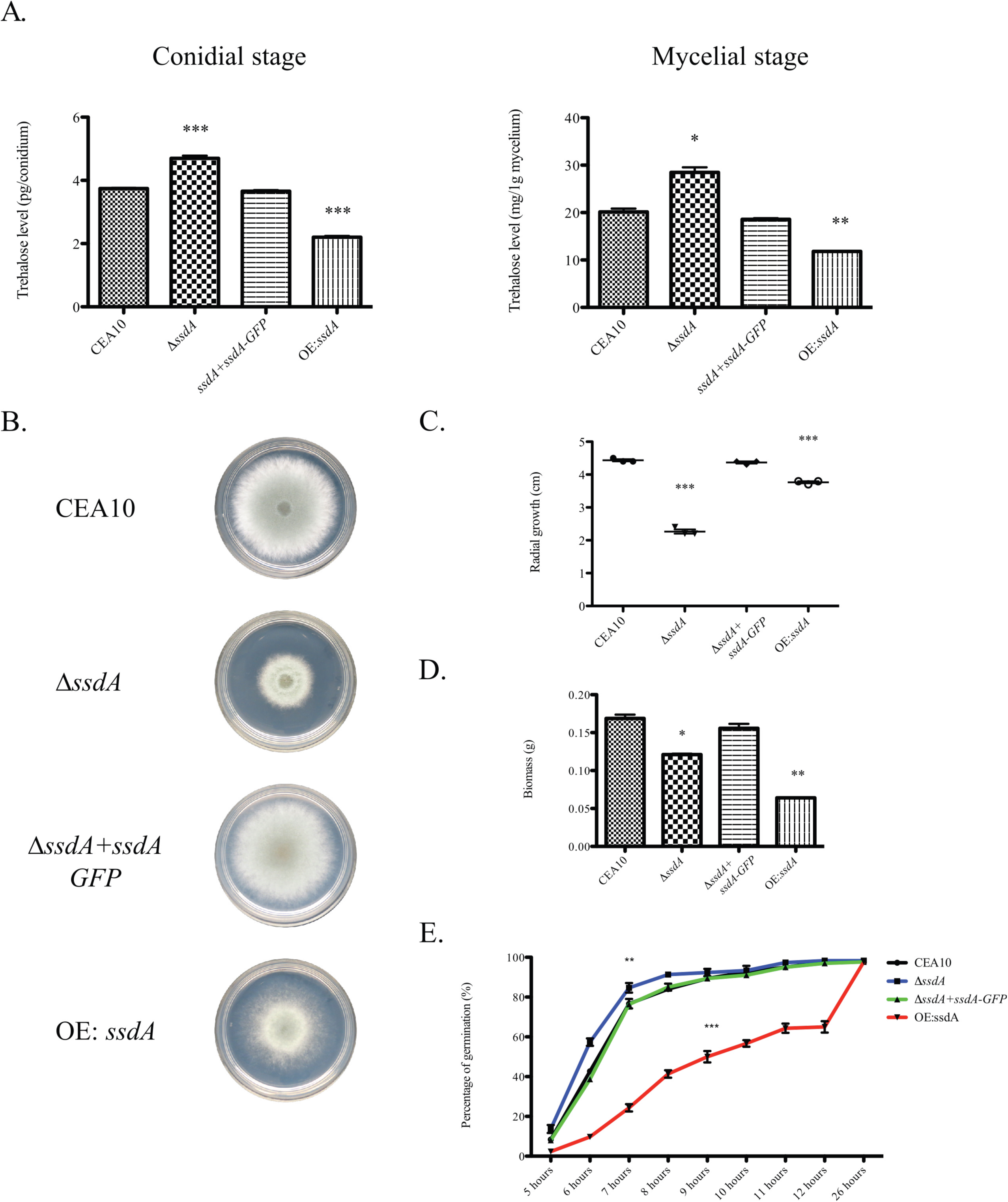
Alteration of *ssdA* expression affects trehalose production, hyphal growth, and conidia germination. (A) Trehalose assays were performed to measure trehalose at both the conidial and mycelial stages using a glucose oxidase assay. Data represented as mean +/− SE of three biological replicates. For the conidial stage, (***) indicates *P* value = 0.0004, unpaired two-tailed Student’s *t*-test Δ*ssdA* to the wild type; (***) indicates *P* value < 0.0001, unpaired two-tailed Student’s *t*-test compared OE:*ssdA* to the wild type. For the mycelial stage, (*) indicates *P* value = 0.0110, unpaired two-tailed Student’s *t*-test Δ*ssdA* to the wild type; (**) indicates *P* value = 0.007, unpaired two-tailed Student’s *t*-test OE:*ssdA* to the wild type. (B, C) Radial growth assays were performed with each strain using GMM at 37°C for 72 hours (B). Images are a representative image of three independent experiments with similar results. The measurement of the radial growth was performed at 72 hours (C). Data represented as mean +/− SE of three biological replicates. (***) indicates *P* value < 0.0001 (unpaired two-tailed Student’s *t*-test compared to the wild-type CEA10). (D) Fungal biomass was measured using 10^8^ spores in 100mL liquid GMM at 37°C for 24 hours. Data represented as mean +/− SE of three biological replicates. (*) indicates *P* value = 0.0133 and (**) indicates *P* value = 0.0023, unpaired two-tailed Student’s *t*-test. (E) Germination assays were utilized using 10^8^ spores in 10mL liquid GMM at 37°C. 500µL of each culture were taken to count for the percentage of germlings at each time point. Data represented as mean +/− SE of three biological replicates. (**) indicates *P* value *<*0.01, unpaired two-tailed Student’s *t*-test, compared the *ssdA* null mutant to the wild type at 6-8 hours. (***) indicates *P*<0.0001, unpaired two-tailed Student’s *t*-test, compared the overexpression strain to the wild type at 5-12 hours.

To determine if *ssdA* plays a role in conidia germination and polarized growth, we measured radial growth of the respective strains’ fungal mycelium on solid medium (**Fig. 1B, C**) and liquid planktonic culture biomass (**Fig. 1D**) in 1% glucose minimal medium (GMM). We observed that on solid GMM both the Δ*ssdA* and OE:*ssdA* strains exhibited decreased mycelial radial growth after a 72-hour incubation at 37°C compared to the wild type and reconstituted strains (Δ*ssdA*+*ssdA:GFP*) (*P*<0.0001 Δ*ssdA* to the wild type and *P=*0.0001 OE:*ssdA* to the wild type) (**Fig. 1B, C**). In planktonic liquid cultures, 24-hour biomass from Δ*ssdA* and OE:*ssdA* cultures grown at 37°C was reduced compared to the wild-type and reconstituted strains (*P*=0.0133 Δ*ssdA* to the wild type and *P*=0.0023 OE:*ssdA* to the wild type) (**Fig. 1D**). Δ*ssdA* conidia germinated faster in the first 6-8 hours (*P*<0.01, Δ*ssdA* to the wild type) while OE:*ssdA* conidia germinated slower during the first 12 hours and then caught up to the wild type at 24 hours (*P*<0.0001, OE:*ssdA* to the wild type) (**Fig. 1E**). The reduced germination observed in OE:*ssdA* conidia is consistent with the reduction in trehalose levels in these fungal cells as Al-Bader, *et al.* observed that depletion of trehalose content in an *A. fumigatus tpsA/tpsB* double null mutant deficient in trehalose delayed conidia germination (39). However, *A. fumigatus* trehalose mutants do not have *in vitro* growth defects when glucose is the primary carbon source and we cannot attribute *ssdA* mutant growth defects to alterations in trehalose levels. Taken together, these results implicate SsdA in trehalose biosynthesis and support a global role for SsdA in fungal fitness when glucose is the sole carbon source.

### SsdA is important for cell wall integrity

*A. fumigatus* trehalose null mutants including Δ*tslA* have altered cell wall integrity and previous research identified a link between yeast Ssd1p and *Neurospora crassa* GUL-1 (SSD1 homolog) function and the cell wall (39, 40, 50, 54). We next utilized the cell wall perturbing agents congo red (1 mg/mL), calcofluor white (CFW, 50 µg/mL), and the echinocandin caspofungin (2 µg/mL) to test the hypothesis that *A. fumigatus* SsdA is important for cell wall integrity. Δ*ssdA* exhibited increased resistance to cell wall perturbing agents while OE:*ssdA* exhibited increased susceptibility, particularly to congo red and calcofluor white (**Fig. 2**).

**Fig 2.**
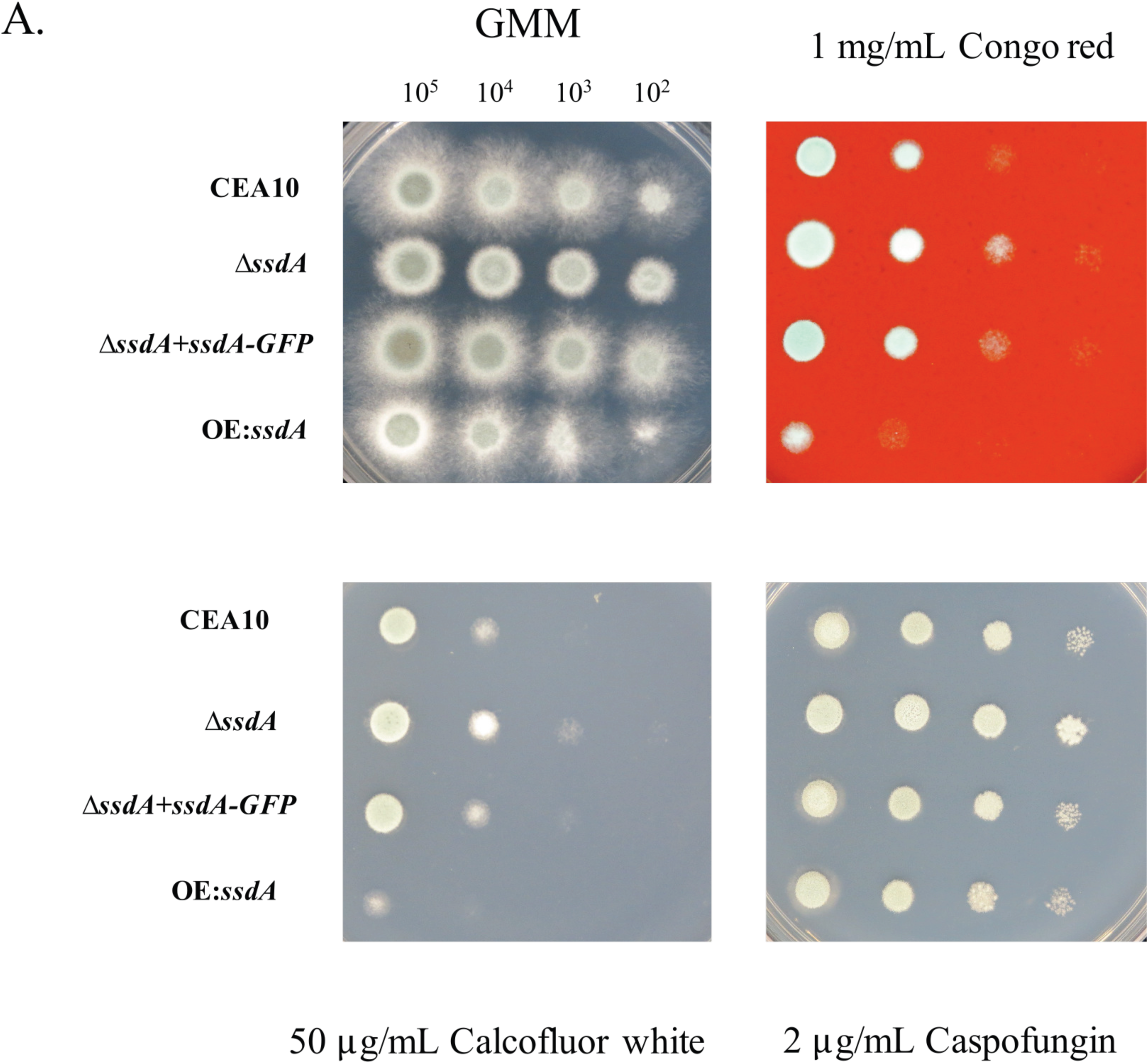
SsdA is important for cell wall integrity. Cell wall perturbing agents, i.e. 1 mg/mL congo red, 50 µg/mL calcofluor white (CFW), and 2 µg/mL caspofungin, were utilized to study cell wall integrity in the respective strains. Cultures were incubated at 37°C for 48 hours. Data are a representative image of three independent experiments all with similar results.

We hypothesized that the altered growth and change in susceptibility to cell wall perturbing agents observed in SsdA mutants comes from altered cell wall composition and/or organization. To initially test this hypothesis, CFW and wheat germ agglutinin (WGA) were used to interrogate total and exposed chitin respectively, while soluble human dectin-1-FC was used to examine β-1,3-glucan exposure. We observed a large decrease in the intensity of CFW and WGA staining of Δ*ssdA* germlings while in contrast germlings of OE:*ssdA* showed increased intensity with these chitin binding molecules (*P=*0.0322, Δ*ssdA* to the wild type, *P*<0.0001, OE:*ssdA* to the wild type) (**Fig. 3A**). For β-1,3-glucan, we observed a decrease in soluble dectin1-FC staining on both the Δ*ssdA* and the OE:*ssdA* germlings suggestive of a decrease in β-1,3-glucan exposure (*P=*0.0389, Δ*ssdA* to the wild type, *P*<0.0001, OE:*ssdA* to the wild type) (**Fig. 3**). While additional quantitative cell wall composition analyses are needed, these data support the hypothesis that SsdA impacts *A. fumigatus* cell wall integrity.

**Fig 3.**
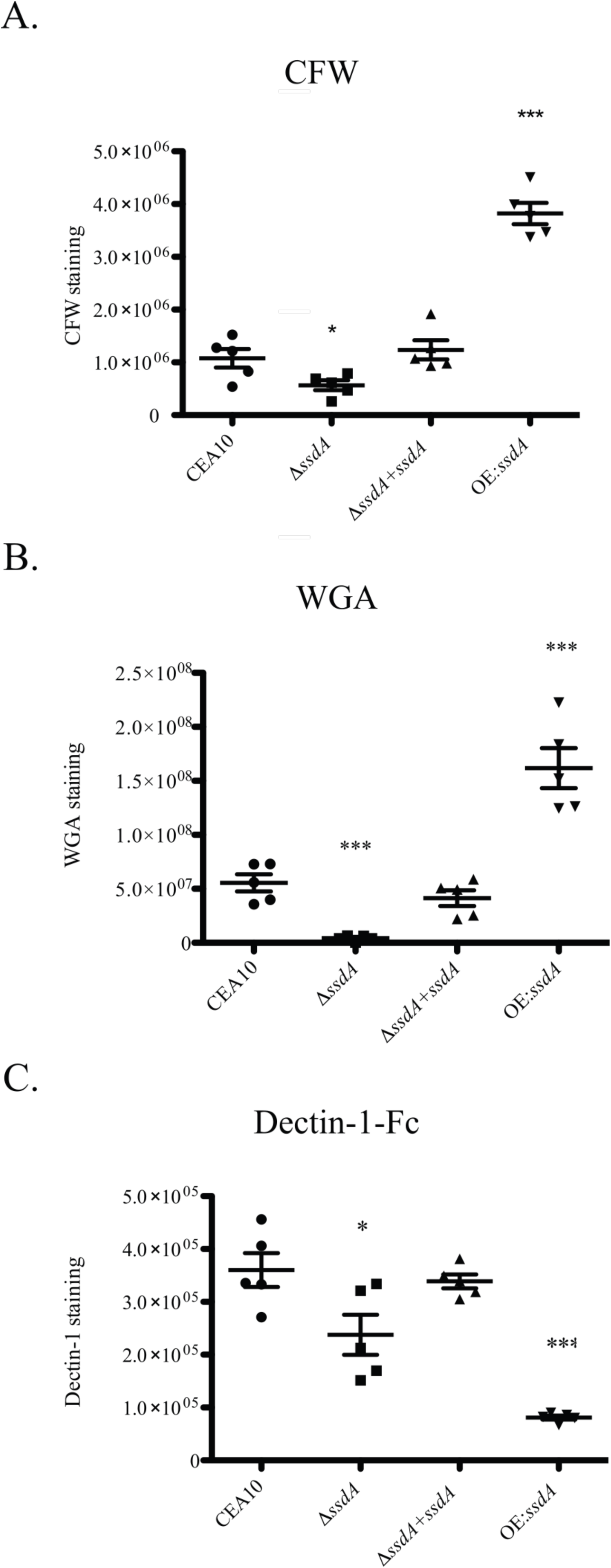
Alteration of *ssdA* expression affects exposure of cell wall PAMPs. Calcofluor white (CFW) staining (A), wheat germ agglutinin (WGA) (B), and soluble dectin-1 (sDectin-1) staining (C) were utilized to observe chitin levels/exposure and the β-glucan exposure on the cell wall of the respective strains. Each strain was cultured to the germling stage under normoxic conditions at 37°C. The corrected total cell fluorescence (CTCF) was calculated. For CFW staining, (*) indicates *P* value = 0.0322, unpaired two-tailed Student’s *t*-test compared Δ*ssdA* to the wild type; (***) indicates *P*<0.0001, unpaired two-tailed Student’s *t*-test compared OE:*ssdA* to the wild type. For WGA staining, (***) indicates *P* value = 0.0002, unpaired two-tailed Student’s *t*-test compared Δ*ssdA* to the wild type; (***) indicates *P* value = 0.0008, unpaired two-tailed Student’s *t*-test compared OE:*ssdA* to the wild type. For sDectin-1 staining, (*) indicates *P* value = 0.0389, unpaired two-tailed Student’s *t*-test compared Δ*ssdA* to the wild type; (***) indicates *P* value *<* 0.0001, unpaired two-tailed Student’s *t*-test compared OE:*ssdA* to the wild type. Data are represented as mean +/− SE of 15 images from three biological replicates. Scale bar 3 µm. AU, Arbitrary Unit.

Given the changes in the cell wall of the Δ*ssdA* and the OE:*ssdA* strains, we next tested their ability to adhere to an abiotic surface. Using the crystal violet adherence assay, we observed no difference in adherence between the wild-type, Δ*ssdA,* and reconstituted strains. However, a striking loss of adherence was observed in the OE:*ssdA* strain (**Fig. 4A**). To investigate this adherence difference further, spinning disk confocal microscopy, in combination with the galactosaminogalactan binding FITC labeled soy bean agglutinin (SBA), was utilized. Given the decreased adherence of the overexpression strain, we were surprised that increased expression of *ssdA* resulted in much greater levels of SBA staining, revealing striking differences compared to the wild-type and Δ*ssdA* strains (**Fig. 4B**). As SBA binds to oligosaccharides with alpha- or beta-linked N-acetylgalactosamine and, to a lesser extent, galactose residues, we tested whether mRNA levels of the UDP-glucose 4-epimerase involved in galactosaminogalactan biosynthesis were altered in the *ssdA* mutant strains (55). No significant difference in *uge3* mRNA levels were observed under the conditions examined, suggesting a role for *ssdA* in post-transcriptional regulation of the galactosaminogalactan polysaccharide (**Fig. 4C**). Taken together, these data suggest that *ssdA* expression levels impact fungal adherence.

**Fig 4.**
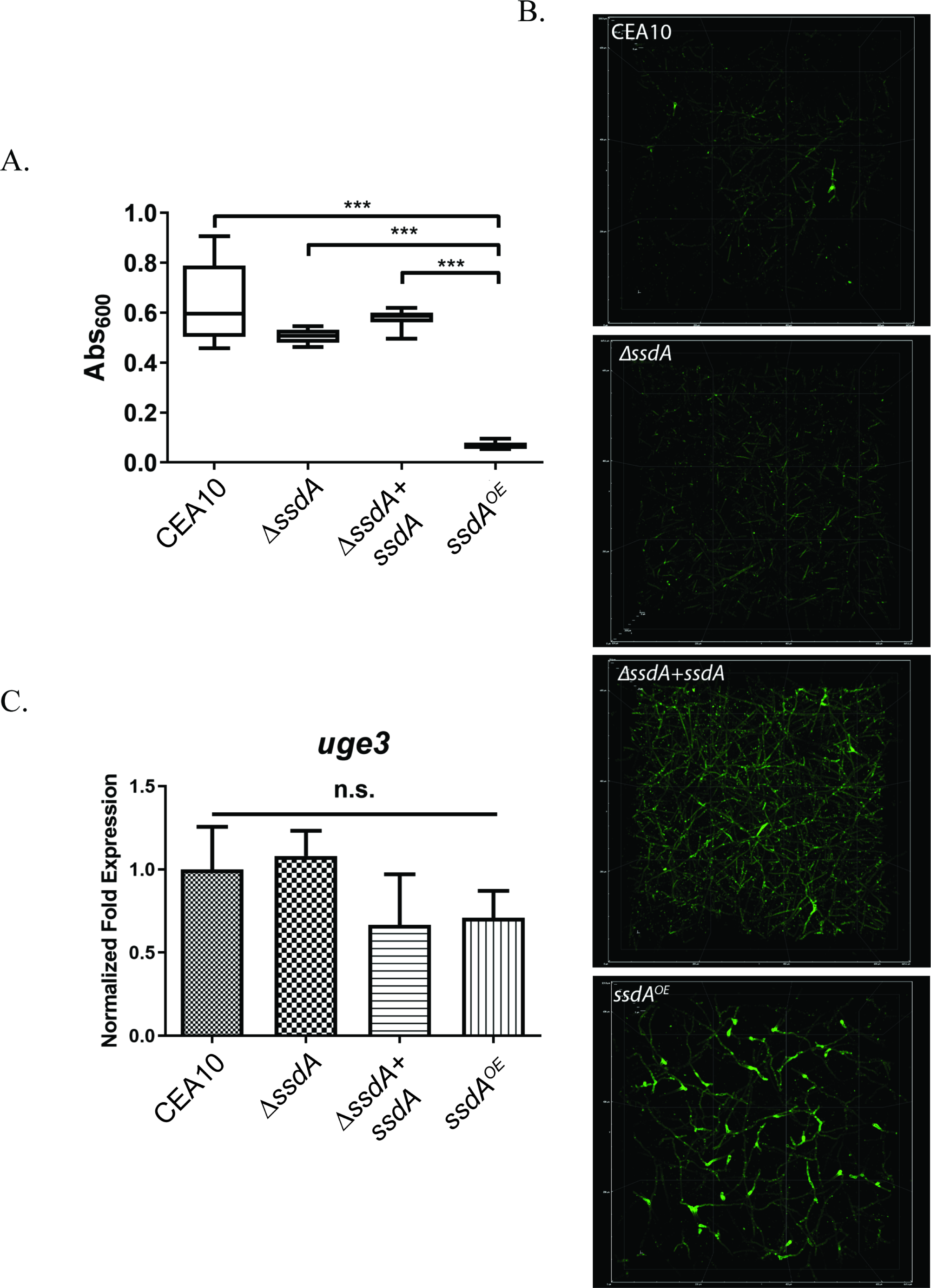
Expression of *ssdA* Impacts Adherence and Biofilm Formation. **(A)**. Biofilms were grown at 37°C for 24 hours in wells of a 96 well plate and the crystal violet adherence assay was performed. Bars represent 6 replicates per strain, and the experiment was repeated 3 times with the same results. (***) indicates *P* value < 0.0001 via One-Way ANOVA with a Tukey post-test. **(B).** Micrographs of 24-hour biofilms stained with FITC-conjugated soybean agglutinin. Images are looking down a Z-stack of the first 300-320µm of the biofilm. Images are representative of 3 biological replicate cultures. **(C).** RNA was obtained from 24-hour biofilm cultures and qRT-PCR was performed for *uge3* mRNA levels. Data were normalized to *tef1* transcript levels. (n.s.) indicates not significant by One-Way ANOVA with a Tukey post-test.

Given the responses of the *ssdA* mutant strains to agents and reagents that inhibit or bind to chitin, a non-radioactive chitin synthase activity assay was next utilized to further define the impact of SsdA levels on the *A. fumigatus* cell wall (56, 57). Consistent with the cell wall immunohistochemistry results, chitin synthase activity in Δ*ssdA* was significantly reduced while in contrast chitin synthase activity in OE:*ssdA* was significantly increased (*P=*0.0029, Δ*csmA* to the wild type; *P=*0.0208, Δ*ssdA* to the wild type; *P*<0.0001, OE:*ssdA* to the wild type) (**Fig. 5A**). We previously observed that activity and localization of the chitin synthase CsmA was perturbed by loss of the trehalose regulatory protein TslA (41). As TslA was found to also physically interact with SsdA, we hypothesized that SsdA levels may also impact CsmA sub-cellular localization.

**Fig 5.**
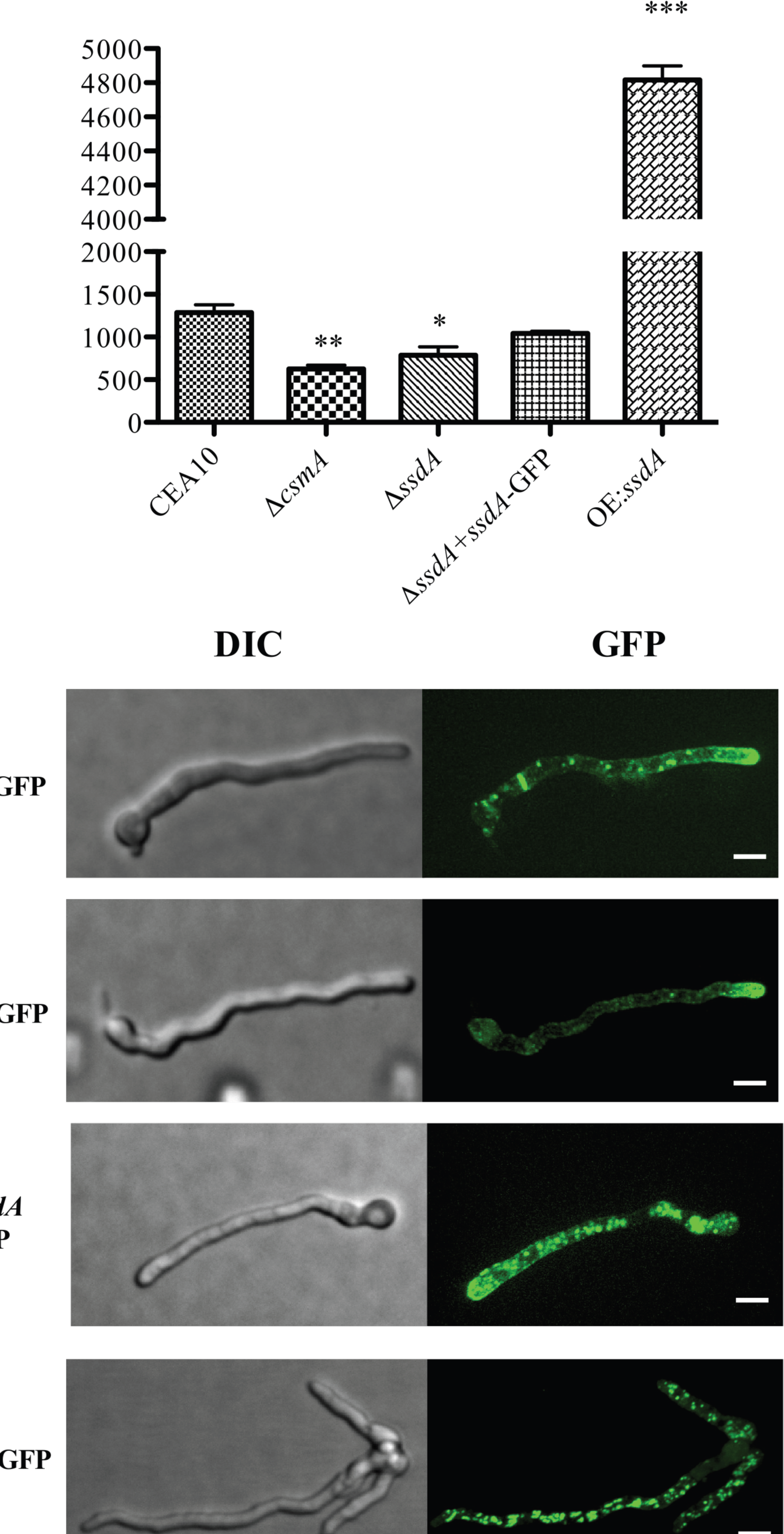
Chitin activity and CsmA localization. **(A)**. 10 µg of membrane proteins were used to perform a non-radioactive chitin synthase activity assay. Each strain was cultured at 30°C for 6 hours and switched to 37°C for 24 hours. (**) indicates *P* value = 0.0029, unpaired two-tailed Student’s *t*-test compared Δ*csmA* to the wild type; (*) indicates *P* value = 0.0208, unpaired two-tailed Student’s *t*-test compared Δ*ssdA* to the wild type; (***) indicates *P value* < 0.0001, unpaired two-tailed Student’s *t*-test compared OE:*ssdA* to the wild type. Data represented as mean +/− SE of three biological replicates. (B) C-terminal GFP-tagged CsmA was generated in the wild type, Δ*ssdA*, *ΔssdA+ssdA,* and OE:*ssdA* backgrounds. Each strain was cultured at 37°C for 12 hours and live-cell imaging was performed under a Quorum Technologies WaveFX Spinning Disk Confocal Microscope (1000X). The images were analyzed using Imaris 8.1.4 software. Data are representative images of 15 images from three biological replicates. Scale bar 3 µm.

To study CsmA sub-cellular localization when SsdA levels are altered, we introduced a C-terminal GFP-tagged *csmA* allele into the respective *ssdA* mutant strains. Using spinning disk confocal microscopy, we observed that alteration of *ssdA* mRNA levels (loss or increase) led to an altered CsmA localization pattern compared to the wild-type and reconstituted strains (**Fig. 5B**). CsmA:GFP puncta observed in Δ*ssdA* are mainly focused at the hyphal tip with a few puncta also localized along the lateral hyphal walls but no visible localization at the conidial septum. In contrast, in OE:*ssdA* CsmA:GFP puncta were dispersed throughout the hyphae with no visible puncta at the hyphal tip or conidial septum (**Fig. 5B**). Intriguingly, this latter result is similar to the diffuse sub-cellular localization of CsmA:GFP in the absence of TslA (41). Taken together, these results suggest SsdA levels affect sub-cellular localization of the chitin synthase CsmA.

### SsdA levels are critical for *Aspergillus fumigatus* virulence

Given the trehalose, cell wall, and biofilm phenotypes associated with alterations in SsdA levels, we hypothesized that SsdA plays an important role in *A. fumigatus* fungal-host interactions. To understand the importance of SsdA in the *A. fumigatus*-host interaction, we first utilized the triamcinolone (steroid) murine model of IPA (58). Strikingly, we observed that overexpression of SsdA significantly decreased *A. fumigatus* virulence compared to the wild type (*P=* 0.0033, OE:*ssdA* to the wild type) (**Fig. 6A**). This reduction in virulence was associated with a large reduction in immune cell infiltrate in the bronchoalveolar lavage fluid (BALs) (*P*=0.0159, OE:*ssdA* to the wild type, Mann-Whitney *t-*test) (**Fig. 6B**). Perhaps correspondingly, we observed a significant reduction in fungal growth within the OE:*ssdA* inoculated lungs compared to other strains (**Fig. 6C**).

**Figure 6.**
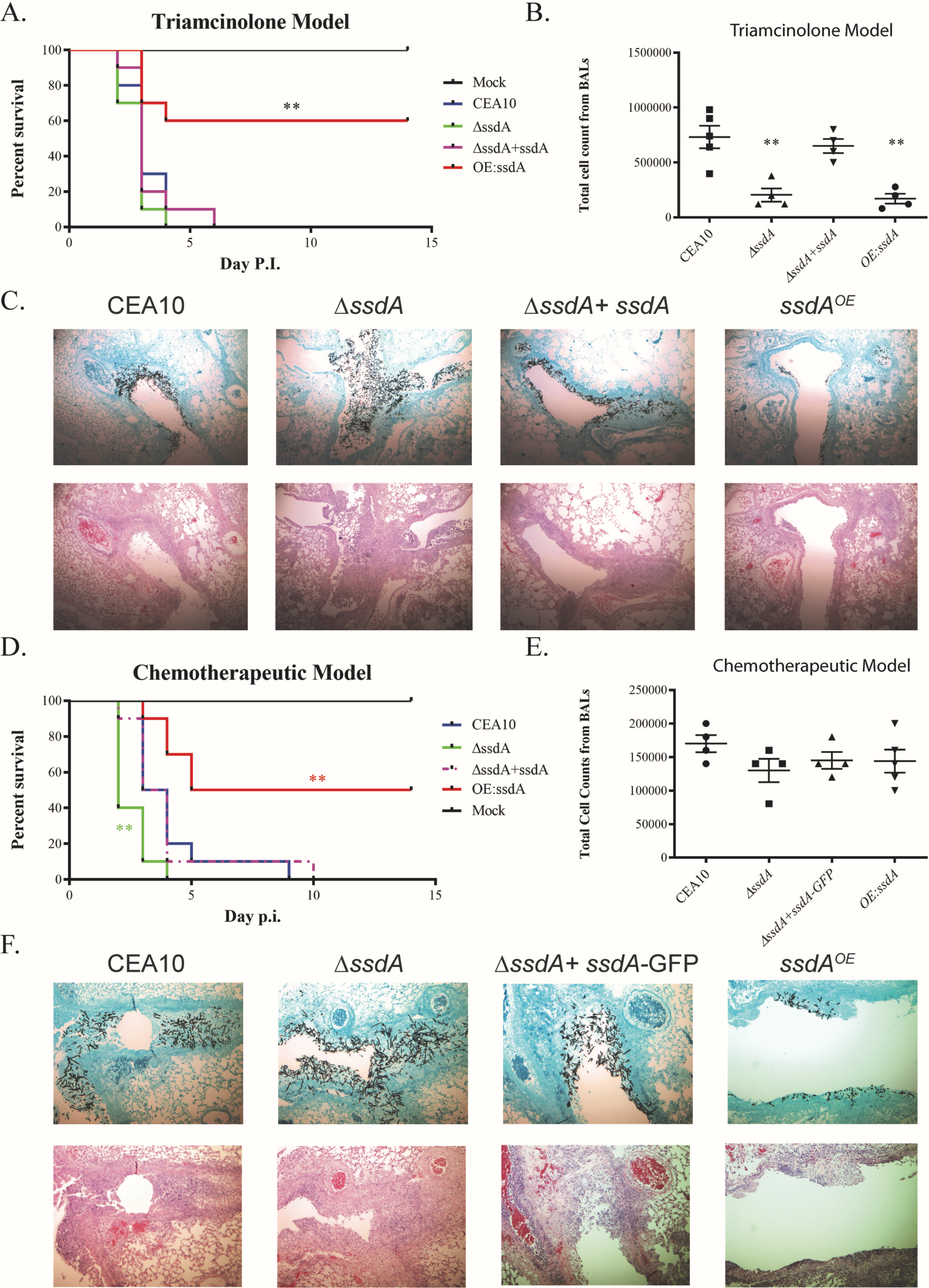
Overexpression of SsdA attenuates virulence in the triamcinolone murine model while both loss of SsdA (Δ*ssdA*) and overexpression of SsdA (OE:*ssdA*) alters immune cell infiltrates. (A) 2×10^6^ conidia of each strain were inoculated via the intranasal route in the corticosteroid IPA murine model. Ten CD1 mice were used in each group. Survival analysis was performed for two weeks. (**) indicates *P* value = 0.0033, Log Rank test compared OE:*ssdA* to the wild type. (B) Δ*ssdA* and OE:*ssdA*-infected BALs had decreased total inflammatory cell infiltrations. Total cell count: (*) indicates *P* value = 0.0159, two-tailed Mann-Whitney *t*-test compared Δ*ssdA* to the wild type; (*) indicates *P* value = 0.0159, two-tailed Mann-Whitney *t*-test compared OE:*ssdA* to the wild type. (C) OE:*ssdA*-infected lungs show less fungal growth and less cell infiltration compared to the wild type. The fungal histology was performed on Day3 to observe fungal growth and inflammatory cell infiltrations. GMS, Gomori-methenamine silver staining; H&E: hematoxylin and eosin staining. Magnification 50×. (D) 1×10^6^ conidia of each strain were inoculated intranasally for the chemotherapeutic murine model and survival analyses were performed for two weeks using ten CD1 mice per group. For Δ*ssdA* mutant, (**) indicates *P* value = 0.005, Log Rank test. For OE:*ssdA*, (**) indicates *P value =* 0.0049, Log Rank test. (B) Δ*ssdA* and OE:*ssdA*-infected BALs had similar total inflammatory cell infiltrations to the wild type BALs. (C) OE:*ssdA*-infected lungs showed less fungal growth compared to the wild type. Histology was performed on Day3 to observe fungal growth and inflammatory cell infiltration. GMS, Gomori-methenamine silver staining; H&E: hematoxylin and eosin staining. Magnification 50x and 100x.

In contrast, complete loss of SsdA did not alter median murine survival time between the wild type and Δ*ssdA* (median survival = 3 days). However, despite the *in vitro* growth defect of Δ*ssdA,* fungal burden observed by histopathology revealed modest increases in Δ*ssdA* fungal burden at day three post inoculation compared to wild-type (**Fig. 6C**). Surprisingly, despite equivalent or increased fungal burden compared to the wild type, a significant reduction in immune cell infiltrate in the bronchoalveolar lavage fluid (BALs) is apparent in animals inoculated with Δ*ssdA* (P=0.0159, Δ*ssdA* to the wild type, Mann-Whitney *t-*test) (**Fig. 6C**). These results support the hypothesis that changes in *ssdA* levels impact the fitness of *A. fumigatus in vivo* and alter host immune responses.

Given the striking Δ*ssdA in vitro* growth defect observed but full virulence (as measured by murine mortality) in the steroid IPA model, we hypothesized that SsdA would be essential for virulence in a leukopenic IPA model with significant immune cell depletion (41). However, surprisingly, and similar to the corticosteroid model, Δ*ssdA* also had persistent if not slightly increased virulence in the leukopenic model (*P = 0.005)* (**Fig. 6D**). Also similar to the steroid model, OE:*ssdA* had significant virulence attenuation compared to the wild type (*P =* 0.0049) (**Fig. 6D**). Median survival of the wild type, Δ*ssdA*, and OE:*ssdA*-inoculated mice was 3.5, 2, and 9.5 days, respectively. Histopathology from this leukopenic model revealed less fungal growth from lungs of OE:*ssdA*-inoculated mice while Δ*ssdA*-inoculated mice, in contrast to the *in vitro* growth phenotype, had substantial invasive hyphal growth compared to the wild type (**Fig. 6F**). In contrast to the steroid model, inflammatory cell infiltrations were the same between *ssdA* mutants and the wild type in this leukopenic model possibly reflecting the significant chemical mediated immune suppression (**Fig. 6E**). These results suggest that increased *ssdA* mRNA levels attenuate *A. fumigatus* virulence likely through fungal fitness defects while loss of SsdA alters the host immune response and modestly increases fungal virulence *in vivo*.

## Discussion

The cell wall of *Aspergillus fumigatus* consists of polysaccharides including chitin, β-glucan, galactosaminogalactan, and others that are critical for fungal fitness in diverse environments including those associated with pathogenesis (29–38). Cell wall homeostasis and integrity are critical for the synthesis of each cell wall component in the face of stress and affect fungal pathogenesis on multiple levels (59). Another carbohydrate produced by fungi, trehalose, is also critical for fungal fitness during environmental stress including pathogenesis (42, 60). Previous research in multiple fungi has revealed an unexpected and ill-defined link between cell wall homeostasis and the biosynthesis of the disaccharide sugar trehalose (39–41). In *A. fumigatus,* a physical interaction between the TslA trehalose biosynthesis regulatory sub-unit and CsmA, a class V chitin synthase, suggested that trehalose biosynthesis proteins have direct roles in coordinating trehalose and fungal cell wall biosynthesis (41). Coordination between these 2 biological processes is logical given that both biosynthetic pathways utilize common carbohydrate metabolic intermediates. Intriguingly in our previous study, TslA was observed to physically interact with a protein (SsdA) that here we define as a homolog of the *S. cerevisiae* translational repressor protein Ssd1p. Alterations in the levels of SsdA in *A. fumigatus* impact both trehalose levels and cell wall integrity. Thus, these data further support the hypothesis that trehalose and cell wall biosynthesis are coordinated and implicate a new potential regulatory protein SsdA in these processes in *A. fumigatus*.

How physical interactions between TslA, SsdA, and CsmA in *A. fumigatus* mediate chitin and trehalose biosynthesis remains unclear. In *S. cerevisiae,* Ssd1p is a unique RNA-binding protein associated with multiple biological processes (43, 44) including stress tolerance, membrane trafficking, cell cycle, posttranslational modifications, mini-chromosome stability, and cell wall integrity (45–47, 61). With regard to a regulatory role in cell wall biosynthesis in yeast, Hogan, *et al.* (2008) showed that mRNA transcripts associated with Ssd1 encoded proteins related to cell-wall biosynthesis, cell-wall remodeling and regulation, cell cycle, and protein trafficking (44). Loss of *Sc*Ssd1p from both human and plant yeast isolates impacted cell wall composition by increasing both chitin and mannan content while decreasing β-1,3-glucan (50). Intriguingly, these results in yeast are opposite to those observed here in *A. fumigatus* where SsdA loss appears to decrease chitin content while overexpression of SsdA increased chitin. However, additional cell wall composition biochemical assays are needed to define the impact of SsdA on cell wall composition.

Ssd1 homologs are also associated with cell wall integrity in the human pathogenic yeast *Cryptococcus neoformans* though a *ssd1* loss of function strain displayed only modest susceptibility to cell wall perturbing agents in this pathogenic yeast (62). However, a Ssd1 homolog in the human pathogenic yeast *Candida albicans* is associated with cell wall integrity and virulence. Increased expression of *CaSSD1* is associated with antimicrobial peptide resistance, while *ssd1* deletion mutants exhibit decreased virulence in an invasive candidiasis murine model (51). Intriguingly, *Ca*Ssd1p physically interacts with *Ca*Cbk1p, an NDR kinase (Nuclear Dbf2-related), which is important for hyphal morphogenesis, the RAM pathway (Regulation of Ace2 and Morphogenesis), polarized growth, cell proliferation, apoptosis, and cell wall biosynthesis (63, 64). *Ca*Ssd1p has nine *Ca*Cbk1p phosphorylation consensus motifs. *Ca*Cbk1p is essential for Ssd1p localization to polarized growth areas (63). Moreover, in the filamentous fungus *Neurospora crassa,* a *gul-1* (Ssd1 homolog) mutant is able to partially suppress the severe fitness defect of a *cot-1* (Cbk1 homolog) temperature sensitive mutant and this is associated with a reduction in transcript levels of cell wall homeostasis genes including chitin synthases and the beta 1,3 glucan synthase *fks1* (54, 65).

The putative *A. fumigatus* Cbk1 homolog (AFUB_068890) is uncharacterized, but the corresponding homolog in *A. nidulans,* CotA, is a conditionally essential gene and it is unclear if it plays a direct role in cell wall or trehalose biosynthesis (66–68). However, loss of *A. nidulans cotA* phenotypes can be suppressed by osmotic stabilization perhaps suggesting an important role for this kinase in cell wall biosynthesis in *Aspergillus* spp (68). Future experiments with *cotA* loss and/or gain of function mutants in *A. fumigatus* may reveal if this important kinase plays a role in chitin synthase regulation and whether this role is mediated by TslA and/or SsdA. Additional domain specific mutations in TslA/SsdA and/or genetic screens may also help reveal the mechanistic relationship(s) behind the TslA-SsdA protein-protein interaction and chitin biosynthesis.

Importantly for human fungal pathogenesis, our results suggest *A. fumigatus* SsdA plays a role virulence. Clinical and plant yeast isolates with null mutations in *Scssd1* have increased virulence in a DBA/2 murine infection model (50). *Scssd1* null mutants induced more pro-inflammatory cytokine production perhaps consistent with alterations in cell wall composition in Ssd1 mutants (50). In the fungal plant pathogens *Colletotrichum lagenarium* and *Magnaporthe oryzae, SSD1* is also important for pathogenesis (52). It was hypothesized that *SSD1* supported plant infection by evading induction of the plant immune response (52). Interestingly, in *A. fumigatus* the loss of SsdA resulted in virulence similar to wild-type strain as measured by murine mortality in both the corticosteroid and leukopenic murine IPA models despite the *in vitro* colony and planktonic growth defects associated with *ssdA* loss. In fact, in both murine models loss of *ssdA* appeared to promote *in vivo* fungal growth but intriguingly reduced the host immune response. In contrast, overexpression of SsdA severely attenuated virulence and we observed significantly less fungal growth in the OE:*ssdA-*inoculated lungs suggesting that loss of virulence in this strain may be due to poor *in vivo* fitness. The extreme adherence defect of OE:*ssdA* may contribute to this loss of *in vivo* fungal burden and virulence, but we cannot rule out other mechanisms impacted by increased *SsdA* levels. For example, the significant delay in conidia *in vitro* germination observed in the OE:*ssdA* strain may also manifest *in vivo* and give the host immune system additional time to clear the fungus. Perhaps consistent with altered cell wall composition and PAMP exposure, both *ssdA* and OE:*ssdA-*inoculated BALF had decreased inflammatory cell infiltration, particularly neutrophils, and how these alterations in the host inflammatory response mediated infection outcomes in the presence and absence of SsdA require further investigation.

In conclusion, we identified a critical role for *A. fumigatus* SsdA in cell wall homeostasis, trehalose production, and virulence. SsdA is involved in regulation of chitin biosynthesis and/or homeostasis in this fungus, however, the mechanisms of this regulation are still unclear and further investigation is needed to fully understand the roles and mechanisms of SsdA in *A. fumigatus* cell wall integrity and fungal-host interactions. While there is a clear conservation of a role for SsdA homologs in cell wall homeostasis in many fungi, these data in *A. fumigatus* provide another example of altered wiring/functions of key master regulatory genes in pathogenic fungi compared to model organisms. It will also be interesting and important to explore the regulation and function of these pathways within fungal species to identify broadly conserved mechanisms for potential therapeutic development.

## Materials and methods

### Fungal strains, media, and growth conditions

*Aspergillus fumigatus* strain CEA17 strain (a uracil auxotroph strain lacking *pyrG* gene) was used to generate the *ssdA* null mutant (69). A *ku80* strain (a uracil auxotroph strain lacking *pyrG* and *akuB* genes) was used to generate S-tagged and Flag-tagged strains for pulldown assays and co-immunoprecipitation experiments (69, 70). Glucose minimal media (GMM) containing 1% glucose were used to grow the mutants along with a wild type, CEA10 (CBS144.89) at 37°C with 5% CO_2_ if not stated otherwise (71). The conidia from each strain were collected in 0.01% Tween-80 after 72-hour incubation at 37°C with 5% CO_2_. Fresh conidia were used in all experiments.

### Strain construction and fungal transformation

Gene replacements and reconstituted strains were generated as previously described (40, 58). PCR and Southern blot were used to confirm the mutant strains (40). Real-time reverse transcriptase PCR was used to confirm expression of the re-introduced gene and overexpressed strain (72). To generate the single-null mutant, *A. parasiticus pyrG* from pJW24 was used as a selectable marker (73). To generate reconstituted strains of single null mutants, we utilized a *ptrA* marker, which is a pyrithiamine resistance gene from *A. oryzae* (74). To generate GFP-tagged strains, we utilized a *hygB* marker, which is a hygromycin B phosphotransferase gene as a hygromycin resistant marker (75). For S-tagged strains, an S-tag coding sequence along with *AfpyrG* was introduced to the C-terminus of proteins of interest, i.e. TslA (76, 77). For co-immunoprecipitation experiments, we introduced GFP-tag with a *hygB* marker into C-terminus of SsdA in the background of C-terminal Flag-tagged CsmA with *pyrG* as a marker (78). In localization experiments, we generated C-terminal GFP-tagged CsmA in both the wild type (CEA17) and the Δ*ssdA* background by using *pyrG* and *ptrA* as a selectable marker, respectively. After the constructs were generated, polyethylene glycol-mediated transformation of fungal protoplasts was performed as previously described (79). For the *ptrA-*marker transformation, we added pyrithiamine hydrobromide (Sigma P0256) into 1.2 M sorbitol (SMM) media at 0.1 mg/L (74). For the *hygB-*marker transformation, we recovered the strains containing the *hygB* marker by adding hygromycin B (Calbichem 400052) into the 0.7% top SMM agar at 150 μg/mL the day after transformation (75).

### Germination assays and Biomass assays

10^8^ conidia in 100mL LGMM of each strain were cultured at 37°C in three biological replicates. 500 μL of each culture was taken to count for germling percentage at indicated time points. For biomass assays, 10^8^ conidia in 100mL LGMM of each strain were cultured for 24 hours at 37°C in three biological replicates. The biomass was collected, lyophilized and dry weight was recorded.

### Cell wall perturbing agents and antifungal agents

Several cell-wall perturbing agents were utilized for cell wall integrity tests: Congo red (CR, Sigma C6277), Calcofluor white (CFW, Fluorescent brightener 28, Sigma F3543), and Caspofungin (CPG, Cancidas, MERCK&CO., INC.). CR, CFW, or CPG were added into GMM plates at final concentrations of 1 mg/mL, 50 μg/mL, and 1 μg/mL, respectively. Dropout assays were performed by plating serial conidial dilutions from 1×10^5^ to 1×10^2^ conidia in a 5-μL drop of each strain. The plates were cultured at 37°C with 5% CO_2_ and the images were taken at 48 hours. This experiment was performed in three biological replicates (40).

### Cell-wall PAMP exposure

Calcofluor white (CFW, 25μg/mL), fluorescein-labeled wheat germ agglutinin (WGA, 5 μg/mL) (Vector labs: FL-1021), and soluble dectin-1 staining were performed as previously described (80, 81). Briefly, each fungal strain was cultured until it reached the germination stage on liquid glucose minimal media. The hyphae were UV irradiated at 6,000 mJ/cm^2^. The micrographs were taken by the Z-stack of the fluorescent microscope, Zeiss HAL 100 (Carl Zeiss Microscopy, LLC, Thornwood, NY, USA) equipped with a Zeiss Axiocam MRm camera. The intensity was analyzed using ImageJ and the corrected total cell fluorescence (CTCF) was calculated (80, 82). Data are represented as mean +/− SE of 15 images from three biological replicates.

### Adherence Assay and Biofilm Microscopy

For the crystal violet adherence assay, 100µL of 10^5 spores per mL in GMM were inoculated into U-bottomed 96-well plates and grown for 24 hours at 37°C. Plates were washed with H_2_O twice, stained with 0.1% (w/v) crystal violet in water for 10 minutes, washed twice more with H_2_O to remove excess stain, and destained with 100% ethanol for 10 minutes. An aliquot of the de-stained supernatants were transferred to a flat-bottomed 96-well plate and Abs_600_ was measured using a plate reader. Results were analyzed using a One-Way ANOVA with a Tukey post-test. For microscopy, 10^5 spores per mL in GMM were grown for 24 hours at 37°C on Mattek dishes (Mattek: P35G-1.5-10-C). Biofilms were stained with 20µg/mL FITC-SBA (Vector Labs: FL-1011) and fixed with 1% paraformaldehyde. Stained biofilms were imaged using a 20X-multi-immersion objective on an Andor W1 Spinning Disk Confocal with a Nikon Eclipse Ti inverted microscope stand with Perfect Focus and equipped with two Andor Zyla cameras and ASI MS-2000 stage. Z-stacks of the first 300-320µm were taken for each sample. Microscopy was performed on three biological replicates per strain.

### RNA Extraction and qRT-PCR

RNA was extracted from 24-hour biofilms grown at 37°C in GMM. Briefly, fungal tissue was flash frozen and bead beat with 2.3mm zirconia/silica beads in 200 µl of TriSure (Bioline: BIO-38032). Homogenized mycelia were brought to a final volume of 1mL and RNA was processed according to manufacturer’s instructions. For qRT-PCR, 5ug of RNA was DNAse treated with Ambion Turbo DNAse (Life Technologies) according to the manufacturer’s instruction. For qRT-PCR DNase treated-RNA was processed as previously described (83). mRNA levels were normalized to *tef1* for all qRT-PCR analyses. Statistical analysis was performed with One-Way ANOVA with Tukey post-test. Error bars indicate standard deviation of the mean (SD).

### Chitin synthase activity assay

10^8^ conidia of each fungal strain were grown at 37°C for 24 hours in 10mL of liquid GMM at 250 rpm. The mycelia were collected to prepare of membrane fractions by a centrifugation at 100,000g for 40 min at 4°C as described before. After that, the nonradioactive chitin synthase activity assay was performed in a 96-well plate as previously described (56, 57).

### Trehalose measurement

Trehalose content in conidia and mycelia was as previously described (40). Briefly, *A. fumigatus* strains were grown on GMM plates at 37°C for 3 days. A total of 2 × 10^8^ conidia were used for the conidial stage of the trehalose assay, and 1 × 10^8^ conidia in 10mL LGMM were cultured overnight for the mycelial stage as described by d’Enfert C and Fontaine (1997) (53). Cell-free extracts were then tested for trehalose levels according to the Glucose Assay Kit protocols (Sigma AGO20). Results from biological triplicate experiments were averaged, standard deviation calculated, and statistical significance determined (*P*< 0.05) with a two-tailed Student’s *t*-test.

### Murine models of invasive pulmonary aspergillosis

CD1 female mice, 6–8 weeks old, were used in the triamcinolone (steroid) or the chemotherapeutic murine model experiments as previously described (40, 41, 58). Mice were obtained from Charles River Laboratories (Raleigh, NC). For survival studies and histopathology, 10 mice per *A. fumigatus* strain (CEA10, Δ*ssdA*, *ΔssdA+ssdA-GFP,* and OE:*ssdA*) were inoculated intranasally with 2×10^6^ conidia in 40 μL of phosphate-buffered saline (PBS) for the triamcinolone model and 1×10^6^ conidia for the chemotherapeutic model, and monitored three times a day. Mice were observed for 14 days after the *A. fumigatus* challenge. Any animals showing distress were immediately humanely sacrificed and recorded as deaths within 24 hrs. No mock inoculated animals perished. Statistical comparison of the associated Kaplan-Meier curves was conducted with log rank tests (84). Lungs from all mice sacrificed at different time points during the experiment were removed for differential cell count and histopathology.

### Histopathology

Three mice in each group (CEA10, Δ*ssdA*, *ΔssdA+ssdA-GFP,* and OE:*ssdA*) were humanely euthanized at day 3 post-inoculation. Lungs were harvested from each group and fixed in 10% formalin before embedding in paraffin. 5μm-thick sections were taken and stained with either H&E (Hematoxylin and Eosin) or GMS (Gomori-Methenamine Silver stain) as previously described (85). The microscopic examination was performed on a Zeiss Axioplan II microscope and engaged imaging system. Images were captured at 50x magnification as indicated in each image.

### Collection and analysis of bronchoalveolar lavage fluid (BALF)

At the indicated time after *A. fumigatus* instillation, mice were euthanized using CO_2_. Bronchoalveolar lavage fluid (BALF) was collected by washing the lungs with 2 mL of PBS containing 0.05M EDTA. BALF was then centrifuged and the supernatant collected and stored at −20°C until analysis. BAL cells were resuspended in 200 µl of PBS and counted on a hemocytometer to determine total cell counts. Cells were then spun onto glass slides using a Thermo Scientific Cytospin4 cytocentrifuge and subsequently stained with a Diff-Quik staining kit (Electron Microscopy Sciences) for differential cell counting (80).

### Ethics statement

This study was carried out in strict accordance with the recommendations in the Guide for the Care and Use of Laboratory Animals of the National Institutes of Health. The animal experimental protocol was approved by the Institutional Animal Care and Use Committee (IACUC) at Dartmouth College (protocol number cram.ra.1).

## Acknowledgments

We thank the members of the microscope facilities in the Department of Biology at Dartmouth College. A.T. thanks Dr. Dawoon Chung for initial training in *A. fumigatus* molecular genetics techniques. A.T. and R.A.C. thank Dr. Jarrod Fortwendel for the chitin synthase activity assay protocol and Thomas Hampton for suggestions on statistical analysis. A.T. also thanks the Department of Microbiology, Faculty of Medicine, Chulalongkorn University, Bangkok, Thailand for fellowship support.

